# “Rotating Frame Relaxation for Magic Angle Spinning Solid State NMR, A Promising Tool for Characterizing Biopolymer Motion”

**DOI:** 10.1101/2022.07.02.498561

**Authors:** Eric Keeler, Ann McDermott

## Abstract

Magic angle spinning NMR rotating frame relaxation measurements provide a powerful experimental strategy to probe biomolecules dynamics, as is illustrated by numerous recent applications. We discuss experimental strategies for this class of experiments, with a particular focus on systems where motion-driven modulation of the chemical shift interaction is the main mechanism for relaxation. We also explore and describe common strategies for interpreting the data sets and extracting motion timescale, activation energy, and angle or order parameters from rotating frame relaxation data. Using model free analysis, and numerical simulations, including time domain treatment, we explore conditions under which it is possible to obtain accurate and precise information about the timescales of motions. Overall, with rapid technical advances in solid state NMR, there is a bright future for this class of studies.

## Studies of Biological Systems

Understanding biopolymer thermal motions is central to the structure-function relationship. Essentially all macromolecular properties of interest from binding thermodynamics to catalysis require fluctuations, and therefore prediction of these functional properties has required knowledge of biopolymer dynamics. Nuclear magnetic resonance, including measurements in solution and solid state, has been a powerful experimental tool for probing biopolymer dynamics on timescales from minutes to picoseconds. Experimental strategies for characterizing dynamics using solid state NMR have been a particularly active area of research recently. ^1^ ^2^

We explore the case of rotating-frame relaxation measurements with magic angle spinning, which have developed into the technique du jour for studying microsecond-to-millisecond motions, encompassing many events of great functional importance such as binding-related rearrangements and allosteric responses.^2^ In these experiments, conformational exchange processes cause large fluctuations in chemical shifts, dipolar couplings and other local anisotropic interactions, since the motions generally change the angle of the functional group with respect to the applied static field. Also, the changes in local environment associated with conformation exchange can lead to additional (typically smaller) changes in shifts and couplings, analogous to the case for exchange detected by solution NMR. Fluctuations in local interactions in turn lead to the decay of nonequilibrium transverse magnetization. In these experiments, magnetization is first prepared by flipping it from its equilibrium position along z to an orthogonal direction, and the decay of this transverse component is measured, typically on the millisecond timescale. In order to suppress dephasing or other spin coherent effects, the experiments generally involve a spin-lock, which has the desired effect of refocusing evolution of the isotropic chemical shift; in this way, in absence of polymer dynamics the magnetization would persist for very long times. For proteins, these measurements are made more incisive yet by joining this relaxation experiment to additional chemical shift dimensions so that the magnetization from individual nuclei can be spectrally resolved, and dynamics information is obtained for individual amino acids, in parallel for the whole molecule. Apart from the promise of biophysical insights, measuring rotating frame relaxation (R_1ρ_) relaxation is of practical interest to the spectroscopist. Like transverse relaxation (R_2_), rotating frame relaxation (R_1ρ_) governs the detection efficiency in a number of pulse sequences that involve spin-locks to effect magnetization transfer, or to simplify (edit or decouple) the spectrum.

Rotating-frame relaxation measurements for solids, where the use of rotating frame relaxation is parlayed into a description of motions using two parameters, a correlation time, τ_c_, and an order parameter, S, that represents the amplitude of the motion (see below for a more detailed description of how the order parameter is defined in NMR studies), are broadly reminiscent of the analogous measurements carried out in solution, which is a popular and mature experimental strategy for characterizing dynamics of soluble proteins.^3,4^ Due to advances in theory, software, hardware, pulse sequences, and sample preparations, solid state NMR (SSNMR) has seen continued growth as a tool for biopolymer studies leading to highly resolved multidimensional correlation spectra. Naturally this has led to the exploration of analogous rotating frame relaxation experiments in solid state NMR. Many parallels exist in the applications of these methods for solids and solution. As for the case of solution NMR, rotating frame relaxation measurements are in some cases combined with other relaxation techniques, in particular R_1_, to probe a larger range of timescales, from picoseconds to microseconds, and elucidate multiple motions that contribute to the measurements. In both cases, extension of these measurements to include multiple experimental temperature points allows for the extraction of an energy of activation for the analyzed processes. For both there is a fertile interdisciplinary opportunity when experimental NMR studies are combined with computational molecular dynamics.

There are also numerous distinctions between solution and solid state NMR relaxation measurements. Solid state NMR R_1ρ_ relaxation measurements are expected to have a somewhat more extended timescale of sensitivity; this is because for solid state NMR the changes in local field that drive relaxation tend to be larger, since they include the anisotropic contribution mentioned above. Also, the combination of the spin-lock with magic angle spinning during the relaxation period leads to a number of complexities and opportunities that exist for solid state NMR but not solution NMR, which are explored in this article.

Perhaps the most important distinction between solution and solid state NMR dynamics measurements is the range of applicability. Without a requirement for crystallization or limitations for molecular weight, solid state NMR is applied in situations where many traditional methods would not be applicable, for example where limited molecular tumbling of the molecule precludes the study by solution state NMR. Figure 1 highlights the breadth of the biological systems, from crystalline globular proteins to amyloid fibrils and oligomeric systems to complex membrane proteins, that have been studied using solid state R_1ρ_ relaxation. Early solid state R_1ρ_ relaxation studies explored well behaved globular proteins, in a microcrystalline state e.g., ubiquitin and GB_1_. Additional studies have further applied the use of ^15^N R_1ρ_ relaxation to examine dynamics in amyloid fibrils and membrane proteins in lipid bilayers in addition to the crystalline protein applications.

**Figure 1:**
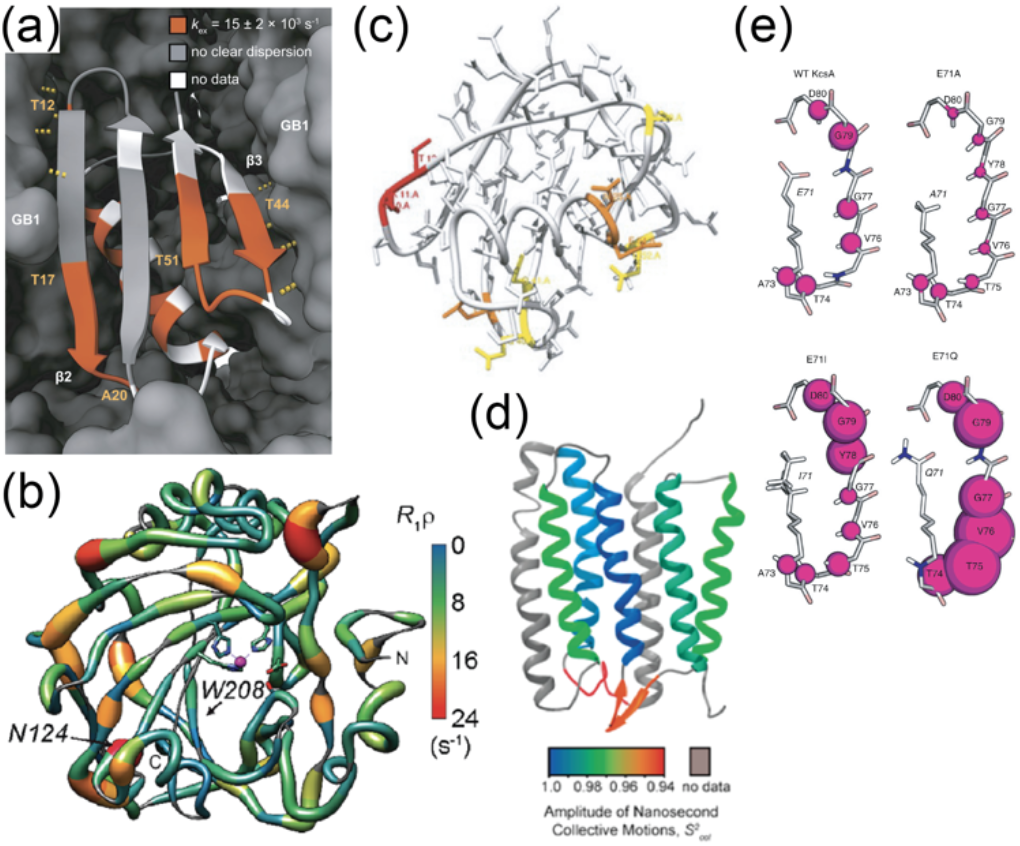
Ribbon diagrams demonstrating the residue specific results of solid state NMR rotating frame relaxation studies of various globular proteins (a) GB1, ^9^ (b) hCAII^10^ and (c) UBQ,^11^ and membrane proteins (d) ASR,^12^ and (e) KcsA.^13^

Table 1 emphasizes that there are many formats used for solid state rotating frame relaxation experiments. For example, the various nuclei (^1^H, ^15^N, or ^13^C relaxation) have different opportunities and advantages for elucidating motions. One popular format for solid state R_1ρ_ measurements of biopolymers involves amidic ^1^H detected experiments^5^ which is an aspect of solid state NMR that has had many advances recently. Commonly these experiments involve recording the relaxation of the attached ^15^N, for which angular fluctuations of the amide functional group would be expected to modulate both the anisotropic chemical shift of the amidic ^15^N and the dipolar coupling to the directly attached proton. Some studies additionally probe the relaxation of the detected ^1^H. These studies are carried out at relatively high applied magnetic field strength (> 700 MHz), and use fast magic-angle spinning (*v*_r_ ≥ 50 kHz), and often also involves deuteration of the protein sample. With these conditions, protons can be detected with excellent sensitivity. Also, complicating effects from the evolution of spin-spin couplings and other kinds of coherent evolution are attenuated with the use of faster MAS. While some fast MAS equipment is commercially produced, is not yet very broadly available, and applicable methods and best practices are still evolving.

**Table 1.** Some applications of rotating frame relaxation for detecting protein motion by SSNMR.

| Protein | Nuclei |  |  | Measurement |  | # of points | MF or Bloch<br>McConnell (BM)<br>component 1 | MF component 2 /3 |
| --- | --- | --- | --- | --- | --- | --- | --- | --- |
|  | <sup>15</sup>N | <sup>13</sup>C | <sup>1</sup>H | R<sub>1ρ</sub> | R<sub>1</sub> |  | τ<sub>c</sub> (p, S<sup>2</sup>) | τ<sub>c</sub> (and p, S<sup>2</sup>) |
| ASR<sup>12</sup> | X |  |  | X | X | 4-5 | sub μs (S<sup>2</sup> ~ 0.89 – 0.99) | Sub ns (S<sup>2</sup> ~ 0.89 – 0.99) |
| GB1<sup>9</sup> | X |  |  | X |  | 10+ | 67 μs |  |
| UBQ<sup>11</sup> | X |  |  | X |  | 3, variable<br>ω<sub>r</sub> | 0.5-3 us (S<sup>2</sup> > 0.95) | 20-200 ps (S<sup>2</sup> ~ 0.65 - 0.95) |
| KcsA<sup>13</sup> | X |  |  | X |  | 1 | n.a. |  |
| hCAII<sup>10</sup> | X |  |  | X |  | 1 | n.a. |  |
| <sup>†</sup>SH3<sup>14</sup> | X |  |  | X |  | 1 | n.a. |  |
| UBQ<sup>15</sup> | X |  |  | X |  | 3+ | 340 μs (p = 9.3%) - BM |  |
| ASR<sup>16</sup> | X |  |  | X |  | 4 | 15-20 ns (S<sup>2</sup> ~ 0.75-0.999) |  |
| GB1<sup>17</sup> | X |  |  | X | X | 3+ | 80 μs (S<sup>2</sup> > 0.98) |  |
| GB1<sup>18</sup> | X | X |  | X | X | 2 | 50 ns (S<sup>2</sup> ~ 0.9-0.99) | 10-100 ps (S<sup>2</sup> ~ 0.6-0.9) |
| HET-s<sup>6</sup> | X | X |  | X | X | 3+ | <sup>15</sup>N 14.7 μs (S<sup>2</sup> = 0.8-0.9) / <sup>13</sup>C 8.7 μs (S<sup>2</sup> > 0.98) | ~10<sup>-8</sup> (S<sup>2</sup> = 0.7-0.99) and 10<sup>-10</sup> s (S<sup>2</sup> = 0.8-0.99) |
| KcsA<sup>19</sup> | X |  |  | X |  | 1 | n.a. |  |
| KpOmpA<sup>20</sup> | X |  |  | X | X | 1 | n.a. |  |
| UBQ<sup>21</sup> | X |  |  | X |  | 3+ | ~10-99 μs (S<sup>2</sup> > 0.9) |  |
| UBQ<sup>22</sup> | X |  | X | X |  | 3 | n.a. |  |
| Y145Stop Prion protein<sup>23</sup> | X |  |  | X |  | 3+ | 1 ms (pop 2%) | 100-300 us (population 5-15%) |
| TET2 enzyme<sup>8</sup> |  | X |  | X |  | 3+ | 6-55 ns (ortho-C, MF, S<sup>2</sup>=0.4375) | 85-110 μs (para-C, BM) |
| <sup>†</sup>Y145Stop Prion protein<sup>7</sup> | X | X |  | X |  | 1 | n.a. |  |
| <sup>†</sup>SH3<sup>24</sup> | X |  |  | X |  | 2 | 5-70 ns (S<sup>2</sup> ~ 0.65-0.99) |  |
| UBQ<sup>25</sup> | X |  |  | X |  | 1 | n.a. |  |
| hCAII<sup>26</sup> | X |  |  | X |  | 7-10+ | 10-800 μs (S<sup>2</sup> > 0.9) |  |
| <sup>†</sup>SH3<sup>27</sup> | X |  | X | X | X | 3+ | <sup>15</sup>N : 5-360 μs (S<sup>2</sup> > 0.925) / <sup>1</sup>H : 63 μs (S<sup>2</sup> ~ 0.4-0.8) |  |
| NS4B<sup>28</sup> | X |  | X | X |  | 1 | n.a. |  |
| GB1<sup>29</sup> | X | X |  | X |  | 1 | n.a. |  |

<sup>†</sup>Studies that include rotating frame relaxation acquired with moderate MAS spinning (≤ 25 kHz)

A handful of other studies have probed the R_1ρ_ relaxation of ^13^C in biopolymers, extending the range of functional groups in the backbone and sidechain that can be probed. ^13^C relaxation can be driven by fluctuations in the chemical shift of the carbon (which exhibit anisotropies up to 100 ppm) as well as dipolar couplings to directly attached protons, with small additional contributions from other directly bonded spins (^13^C, ^15^N or other). Promising studies of C-alpha,^67^ carbonyl,^2^ and aromatic^8^ carbons in protein systems are among the examples listed in Table 1. The use of specific isotopic enrichment schemes to isolate the ^13^C sites and eliminate strong homonuclear couplings is employed in some cases. Overall, there appears to be many aspects of these measurements yet to explore.

## Experimental Approaches: Rotating Frame Pulse Sequences

As for solution NMR, measuring the rotating frame relaxation of solid biological samples is typically achieved by applying a spin-lock on the nucleus of interest, and monitoring the magnetization decay constant, i.e. the reduction in amplitude of a particular peak in a spectrum with respect to spin-lock duration, as illustrated in Figure 2(a,b), and typically this decay is fit to an exponential, to obtain the rotating frame relaxation rate constant, denoted as R_1ρ_. The general and popular spin-lock pulse sequence element is relatively simple, wherein the spin-lock is achieved simply by simple continuous wave (CW) irradiation, herein referred as a traditional spin-lock (TradSL). A broad range of motions cause readily detected elevations in the rotating frame relaxation rate, changes of about 10 s^-1^ or greater for a modestly anisotropic site, making this experiment an excellent choice for detecting motion on the microsecond timescale. As illustrated in Figure 2, motions that are expected to make marked contributions include motions on timescales from 10^4^ to 10^7^ s^-1^, that involve angles of roughly 10 degrees or more (such that the plot’s abscissa, the order parameter squared, S^2^, described below, which in general has a value 1.0 for rigid sites and 0 for entirely isotropic sites, is less than ~ 0.98).

**Figure 2:**
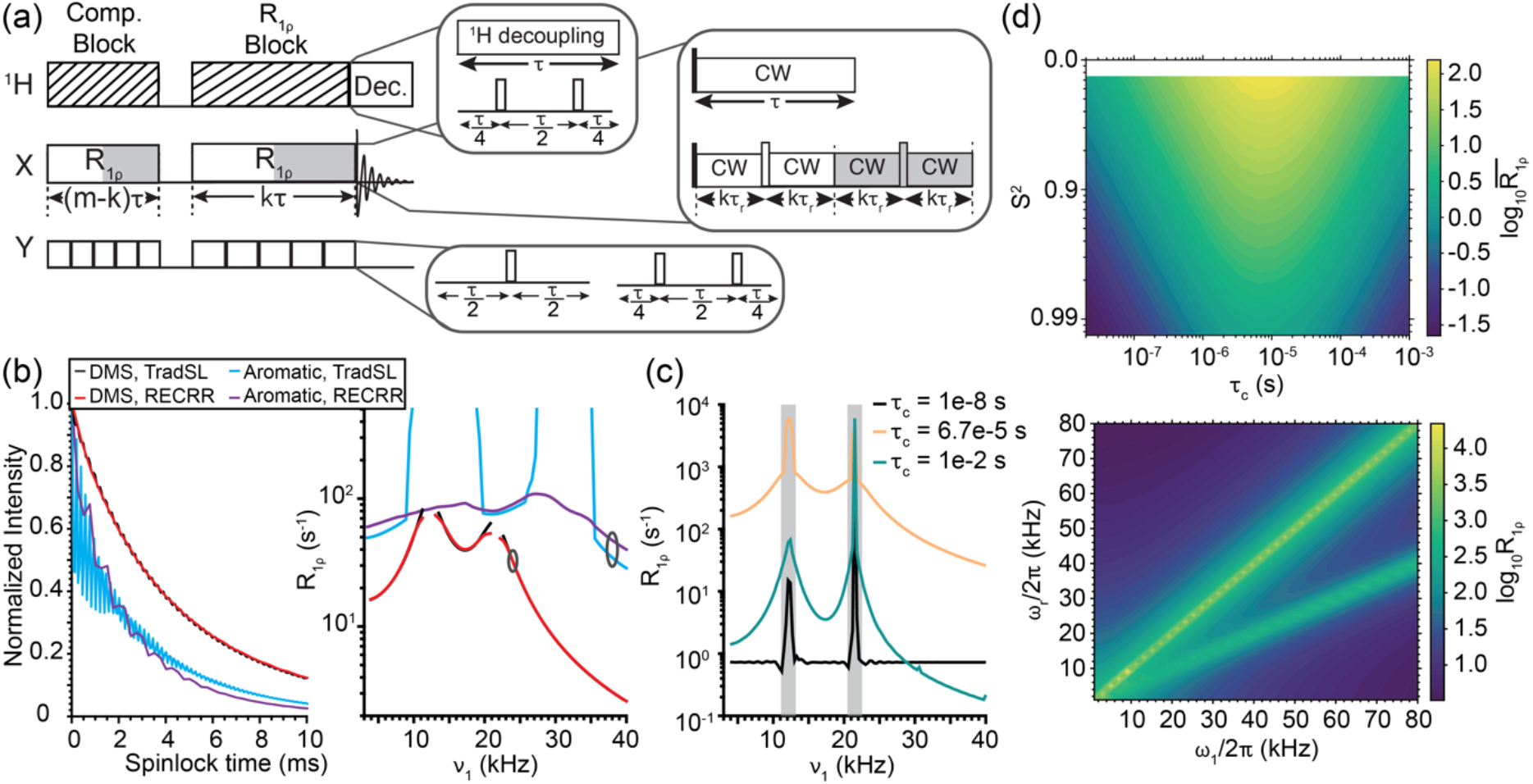
**(a)** Rotating frame relaxation pulse sequences showing the various types of sequences are used on each channel during the spin-lock. **(b)** Numerically simulated time decays for traditional spin-lock (black and cyan) and RECRR (red and purple) experiments on two different systems, DMS (black and red) and an aromatic carbon (cyan and purple) at *v*_1_ = 2.4 *v*_r_ (right). Relaxation dispersion curves for these systems and experiments (right) obtained by fitting the decay of the signal to an exponential function. The gray circle indicates the point depicted on the left. ^13^C sites with highly anisotropic shielding (such as carbonyls and aromatic groups) typically exhibit elevated relaxation in comparison with sites with smaller anisotropy (such as methyl groups). Sites with elevated anisotropy (in comparison with the spinning frequency) also exhibit coherent dephasing, as illustrated in (2b cyan curve). The RECRR sequence refocuses coherent evolution of the anisotropic shift, allowing the relaxation decay to be recorded with greater fidelity (2b purple curve). Simulations of DMS are performed at a static magnetic field of 400 MHz with a spinning frequency of 10 kHz with δ = −36.77 ppm, η = 0, τ_c_ = 67 μs, and β = 109° (S^2^ = 0.3295) and simulations of aromatic carbon are performed at a static magnetic field of 750 MHz with a spinning frequency of 16 kHz, with δ = 175 ppm, η = 0, τ_c_ = 700 μs, and β = 120° (S^2^ = 0.4375). R_1ρ_ rate constants were extracted by fitting the time decay to a single exponential function. **(c)** Numerically simulated time decays of DMS (parameters excepting τ_c_ are the same as **(b)**) demonstrating the difference in the R_1ρ_ dispersion curves between fast limit (τ_c_ = 10 ns), intermediate (τ_c_ = 67 μs), and slow limit (τ_c_ = 10 ms) timescales. **(d)** Contour plots of the extracted R_1ρ_ rate constant for numerical simulations of DMS with respect to the order parameter (S^2^) and the correlation time (τ_c_) (top) and with respect to the applied spin-lock field (ω_1_/2π) and the spinning frequency (ω_r_/2π) (bottom). DMS simulations for the τ_c_ vs S^2^ plot were performed with the following parameters: ω_oH_/2π = 400 MHz, *V*_r_ = 10 kHz, *v*_1_ = 2.55, 3.15, 3.55, 3.85, 4.2 *v*_r_, S^2^ = 0.25 – 1.0, δ = −36.77 ppm, η = 0, τ_c_ = 1e-8 – 1e-3 s with the average rate constant, 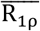, evaluated across the *v*_1_ dispersion curve. The DMS simulations for the ω_1_/2π vs ω_r_/2π plot were performed with the following parameters: ω_oH_/2π = 400 MHz, *v*_r_ = 1-80 kHz, *v*_1_ = 1-80 kHz, β = 109° (S^2^ = 0.3295), δ = −36.77 ppm, η = 0, τ_c_ = 67 μs.

Given the broad range of motions that can result in relaxation, is it possible to discern the specific timescale? Notably, the plot of magnetization decay rate with respect to the motion’s correlation time is “double-sided” (Figure 2(c, top), sloping to smaller rates of decay with an increase in rate for fast limit motions, and sloping to smaller rates of decay with a decrease in correlation times for slow limit motions. The consequence of the “double-sided” dependence of rate with respect to correlation time is that comparative experiments that probe two temperatures or two closely related samples are ambiguous to interpret – an increase in relaxation could be owing to either increased rate constant (it the rate is slow limit) or a decrease in rate constant (if the rate is fast limit), absent additional information or assumptions. In fact, the increase could also be due to increase in angle or amplitude of the motion, as represented by the order parameter S, without change of timescale. Identification of the exchange rate for the motion driving relaxation requires can be done can by repeating the relaxation measurement at various spin-lock field strengths; the plot of rotating frame relaxation rates as a function of applied field strength is referred to as a dispersion curve. The choice of spin-lock strength is critical; since it acts to refocus chemical shift evolution, the effects of exchange on timescales much slower than the refocusing time associated with the spin-lock strength are effectively suppressed. In practice, then, this strength is varied to generate a dispersion in the rate of decay that can be used to identify the timescale of the motion (“dispersion curve” analysis. We use the term “fast limit” which is defined by comparison of the rate constant k_ex_ to other characteristic frequencies involved in the experiment: k_ex_ >> ω_1_ + ω_r_ where ω_1_ is the applied field frequency and ω_r_ is the magic angle spinning frequency. As illustrated in figure 2, dispersion curves are flat for fast limit motion and strongly sloped when the motion is intermediate or slow exchange.^30^ Both the sample spinning and the applied field can cause refocusing of chemical shift evolution of the spin, and both are variable and of the order 1-100 kHz (or mid-microsecond timescale). If spinning or pulse induced refocusing occurs on a timescale much faster than the motion, the effect of the motion is largely quenched. Therefore, slow motions are implicated if fast vs. slow spinning, or strong vs. weak applied fields cause marked change in the relaxation rate. In summary, dispersion curves can provide a ready signature for millisecond vs nanosecond motions, and even supply a potential approach for quantitative determination of timescale. Magic angle spinning introduces features in the dispersion that are unfamiliar from solution NMR curve. Pronounced elevated features are seen at the sample rotation frequency and twice this frequency, which have mixed contributions from unwanted coherent evolution and from the fact that the motion contributions for slower motions are maximal at these frequencies; these effects are discussed more later in this article.

The dispersion curve, i.e. repeating the experiment with a minimum of two different spin-locking fields, solves this problem in principle, and is very incisive to provide at least a qualitative indication of timescale regime, a fact that has been exploited also for solution NMR.

Measurements in solids have added experimental and theoretical complexity related to the sample rotation used to achieve high resolution solid state NMR spectra. The pulse used to refocus the magnetization may lead to more complex effects than for the solution experiment, and does not effectively suppress evolution of dipolar couplings and chemical shift anisotropy. For example, magic angle spinning has the effect of refocusing (or averaging) the evolution of the chemical shielding anisotropy. At first blush, this would seem to be powerful, to employ two independent measures, spinning and the spin-lock, to refocus coherent evolution. However, they interact in a coherent way and, if the spin-lock field strength ω_1_ is matched to the spinning frequency ω_r_, the CSA does not refocus after each rotor period but rather undergoes significant net evolution, an effect referred to as “recoupling” or “reintroduction of the tensor”.^31^ Coherent evolution (CE), defined for the purposes of this article as the evolution of coherent spin interactions during the application of a spin-lock, is therefore expected at these conditions, variously called CSA conditions or R^3^ conditions, where ω_1_ = nω_r_.

The requirement that there be no coherent evolution has been a bottleneck for some SSNMR R_1ρ_ studies in practice, because avoiding these specific “recoupling conditions” and others involving dipolar couplings is not as simple as one might initially think. The matching conditions are expected to unavoidably have a width of the order of the span of the anisotropy and so render many portions of the dispersion curve unsuited to relaxation analysis. Moreover, experimental artefacts such as spatial inhomogeneity in the RF field can lead to additional apparent broadening of these conditions in practice. The need to avoid conditions by a large margin in order to avoid strong coherent evolution is demonstrated in Figure 2(c, bottom). It requires in practice that the span of the CSA be small compared to the spinning frequency ω_r_, that isotopic dilution be used (e.g. by deuteration and use of sparse ^13^C or ^15^N) and that recoupling conditions be avoided in selection of spin-lock strength. Alternatively, special pulse sequences that refocus or defeat the recoupling can be adopted, as discussed below.

One remedy to avoid these conditions and their associated issues is to elevate the spinning frequency ω_r_, and use relatively weak RF fields ω_1_, in particular ensuring that the span of the CSA be small compared to the spinning frequency, and the difference ω_r_ - ω_1_ > δ. Dipolar driven coherent evolution is expected to be attenuated at high frequency spinning as well, removing another potential source of unwanted coherent evolution, though isotopic dilution can also be employed for this purpose. Furthermore, fast spinning has the additional benefit that it is compatible with ^1^H detected experiments and relatively narrow ^1^H lines, which in turn exhibit important advantages in detection sensitivity.^32^ Therefore, some of the applications listed in Table 1 utilize higher spinning frequencies (*v*_r_ ≥ 50 kHz).

Despite this clear advantage, in many cases moderate spinning frequencies have been and continue to be utilized for R_1ρ_ measurements together with direct ^13^C detection. This can motivated by practical considerations, namely the availability of the equipment for this style of experiment, resulting in its being well tested and user friendly. It is also noteworthy that the slower spinning frequency leads to increased sensitivity to intermediate timescale motions for many cases (stronger relaxation rates for the same motion and spin system). A strategy to address unwanted evolution of the CSA at lower spinning frequencies utilizes refocusing pulses during the spin-lock that effectively refocus numerous interactions so that net evolution is suppressed. For example, a pulsed spin-lock sequence element termed the REfocused CSA Rotating-frame Relaxation (RECRR) suppresses evolution of the magnetization under the influence of the anisotropic CSA.^33^

For experiments at lower sample spinning frequencies, some dipole driven unwanted coherent evolution can be avoided by use of sparse isotopic labeling schemes to isolate the spin system and simplify the measurement and its analysis. Also, various decoupling elements have been used on the ^1^H and Y channels with the intention of suppressing unwanted dipolar evolution or to reduce cross-correlated relaxation involving both dipolar and CSA mechanisms. Frequently this has been implemented in the form of one or two π pulses during the spin-lock time. In some cases, high powered CW decoupling is applied on the ^1^H channel while recording X – channel rotating frame relaxation, ideally with an amplitude (*v*_dec_) that avoids magnetization transfer or interference with a pulsed spin-lock element, and ideally without interference between the decoupling and the motion of interest.^34^ In some cases, a compensation block of length related to the rotor period is utilized to ensure that each time point during the decay has an identical total RF load (and associated RF heating) for the experiment. For complex biological systems the relaxation measurement pulse sequence element is inserted into two- or three-dimensional experiments to create a pulse sequence that allows for site-specific measurements of the rotating frame relaxation rates.

The efforts highlighted above point the way to possibile future inventive pulse sequences that suppress a range of unwanted coherent effects, to best preserve magnetization (and result in better sensitivity) and render the relaxation measurements more accurate and precise.

### Strategies for Analysis, Simulation and Determination of Timescale

Various analysis strategies have been attempted for obtaining conformation exchange characteristics from experimental NMR relaxation data. While qualitative indications of mobility can provide insights into biopolymer function and stability, the larger goal of these studies is to obtain quantitative constraints on structure changes and their timescales. A common approach to identifying the angle and timescale is model free analysis (MF). MF analysis involves analytic generalized equations derived from Bloch-Wangness-Redfield theory which were developed initially for biopolymer dynamics in solution.^35,36^ These parsimonious expressions describe expected relaxation rates for a motion or several motions, each characterized by an order parameter and a rate. The MF expressions are user friendly in the sense that they are relatively simple, differentiable and do not require knowledge of the type of motion. This approach has had a significant impact on solution NMR of biopolymers, and there are a number of software packages available^37^. The expressions are expected to be applicable only for a useful but circumscribed regime. The fast limit weak field approximations that are inherent to Redfield theory are assumed, suggesting that these expressions should only be applicable if the correlation time is much faster than the characteristic relaxation time (i.e. faster than the ms timescale) and faster than the inverse of the anisotropic interaction that causes relaxation (here faster than about 20 μs). If multiple motions are included, they should be separable mechanistically. The basic MF expressions also assume that the anisotropy driving the relaxation is uniaxial (cylindrically symmetric). The expressions do not account for any coherent evolution, and so should be used to fit data where coherent evolution has been effectively suppressed.

Modified MF expressions were recently developed for magic ancle spinning (MAS) SSNMR studies^38^. This approach provides relatively simple expressions for a relaxation process driven only by chemical shift anisotropy, as indicated below and a different set of equations for processes driven by fluctuations of the dipolar interaction. The expressions derived for model free prediction of rotating frame relaxation driven by the CSA interaction^38^, and used in this study were as follows.

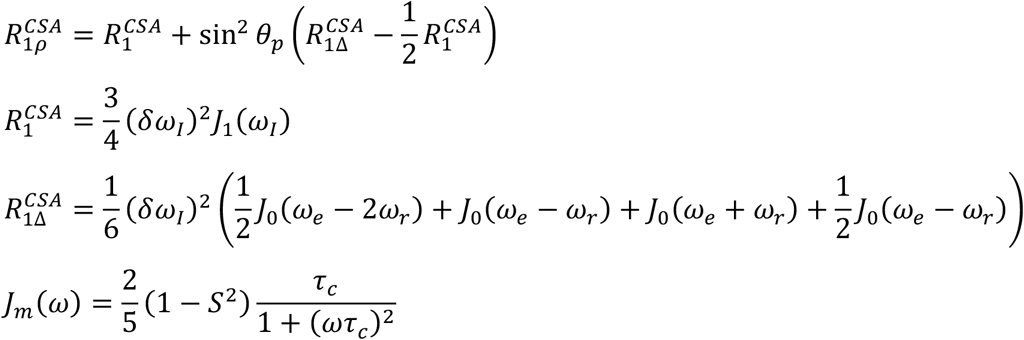

Here CSA refers to the chemical shielding mechanism, R_1Δ_ refers to exchange contributions to the transverse relaxation. J refers to the spectral density associated with the motion, δ_ω1_ refers to the anisotropy of the CSA (σ_zz_ - σ_iso_, the magnitude of which is equal to 2/3 of the full span Ω=σ_11_ - σ_33_, and which is usually approximated to be uniaxial, i.e. the asymmetry parameter η = (σ_yy_ - σ_xx_) /(σ_zz_ - σ_iso_) =0, though that approximation is not generally accurate). In addition, ωe refers to the field strength of the RF excitation, ω_r_ refers to the rotation frequency for magic angle spinning. τ_c_ refers to the correlation function for the motion. S represents the order parameter and has a special definition for NMR studies, characterizing the degree of averaging for the operative tensor due temporal fluctuations in the angle relating the director of the functional group to the applied static magnetic field. As such, it is a tensorial value and a second order function of the angles involved in the motion because the anisotropic interactions generally depend on the angle of the functional group with respect to the applied static magnetic field according to a second Legendre polynomial in angle. On the other hand, for uniaxially symmetric motions and tensors it can be expressed simply as the scalar ratio of the thermally averaged anisotropy to the corresponding static value S = 〈*δ*〉/*δ* (angular brackets represent the time average), which is namely the order parameter as is used in model free analysis. When generating a model free expression to represent a two-site hop Markov model with equal populations and hop angle β, the following expression was used to compute the squared order parameter. ^39 36^

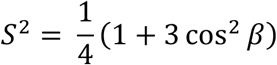

θ refers to the tilt angle for the spin-lock. R_1_ and R_2_ represent other longitudinal and transverse decay rates, respectively. R_1_ and R_2_ are assumed to be driven primarily by other motions that operate on nanosecond to picosecond timescales. R_1_ and R_2_ have also been approximated analogously by a model free expressions, for example if the rate is driven by fluctuations in the CSA, 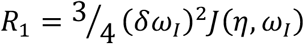 and 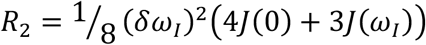. with spectral densities that are assumed to take the same functional form as given above, however due to the differences in the spectral density arguments, would be driven by a process with a shorter time constant than the primary driver for R1Δ.

As is the case for the solution NMR MF analysis, the SSNMR MF expressions are computationally convenient: they are in principle applicable to arbitrary random motion mechanisms described by a generalized order parameter S and a characteristic time constant τ_c_, they are simple expressions, rapid to evaluate, and amenable to differentiation, interpolation, and intuition. As such they have been popular and already been used in the analysis of a number of studies, as illustrated in Table 1. The assumptions listed above for solution NMR applications all apply here as well, namely that no significant coherent evolution should occur, a sometimes vexing point discussed in detail above, and that the interactions should be uniaxial and the motions should be fast compared to the relaxation rates themselves and the interactions that drive them. If these requirements are met, MF can be a handy tool.

If the data do reflect contributions from coherent evolution, or if other assumptions regarding timescale fail, a more flexible and comprehensive approach is possible through numerical simulation of magnetization evolution under the influence of both coherent effects of the Hamiltonian(s) and chemical exchange in the form of an N-site hop mechanism. This approach has been implemented in both SPINEVOLUTION^40^ and SPINACH^41^. If all important interactions are included, this approach is expected to be accurate and flexible, although computationally intensive. Several of the applications listed in Table 1 also employ this kind of strategy, or compare the two strategies.

To illustrate the requirements that allow a unique and correct motion model to be identified from relaxation measurements, we analyzed data for which the motional model is assumed to be known, using both experimental data for a well characterized small molecule, as well as simulated dispersion curves that mimic perfect experimental data. We made simplifying assumptions, for example a single motion, assumed to be a two-site hop, that modulates a uniaxial CSA interaction, with other interactions and causes of dephasing or relaxation omitted. We initially re-examined dimethyl sulfone (DMS), an experimentally forgiving system in the sense that the methyl carbon has a relatively small tensor, and when deuterated the CSA is a dominant term in the Hamiltonian driving relaxation. In a previous study^42^, NMR relaxation measurements on this system at several temperatures were used to extract the activation energy for a two-fold jump motion and associated change in the orientation of the chemical shift anisotropy (CSA) tensor of the methyl carbon. Analysis of these measurements carried out previously used the simulation program SPINEVOLUTION^40^ identified an activation energy (Ea) of between 71 and 75 kJ/mol (Figure 3), in good agreement with previous data and analysis characterizing this activation energies as E_a_ = 63 – 87 kJ/mol. We re-analyzed those data and additional simulated data with an updated workflow, assuming no prior knowledge of rates, and thereby tested a protocol for future analysis of experimental data for new systems. The present analysis of these experimental data utilized the simulation package SPINACH^41^ for numerical simulations as well as other routines in MATLAB including one for model free (e.g. Figures 2 - 7); notebooks with details for reproducing these simulations are available at the website for COMD/NMR at the New York Structural Biology Center (https://comdnmr.nysbc.org).

**Figure 3:**
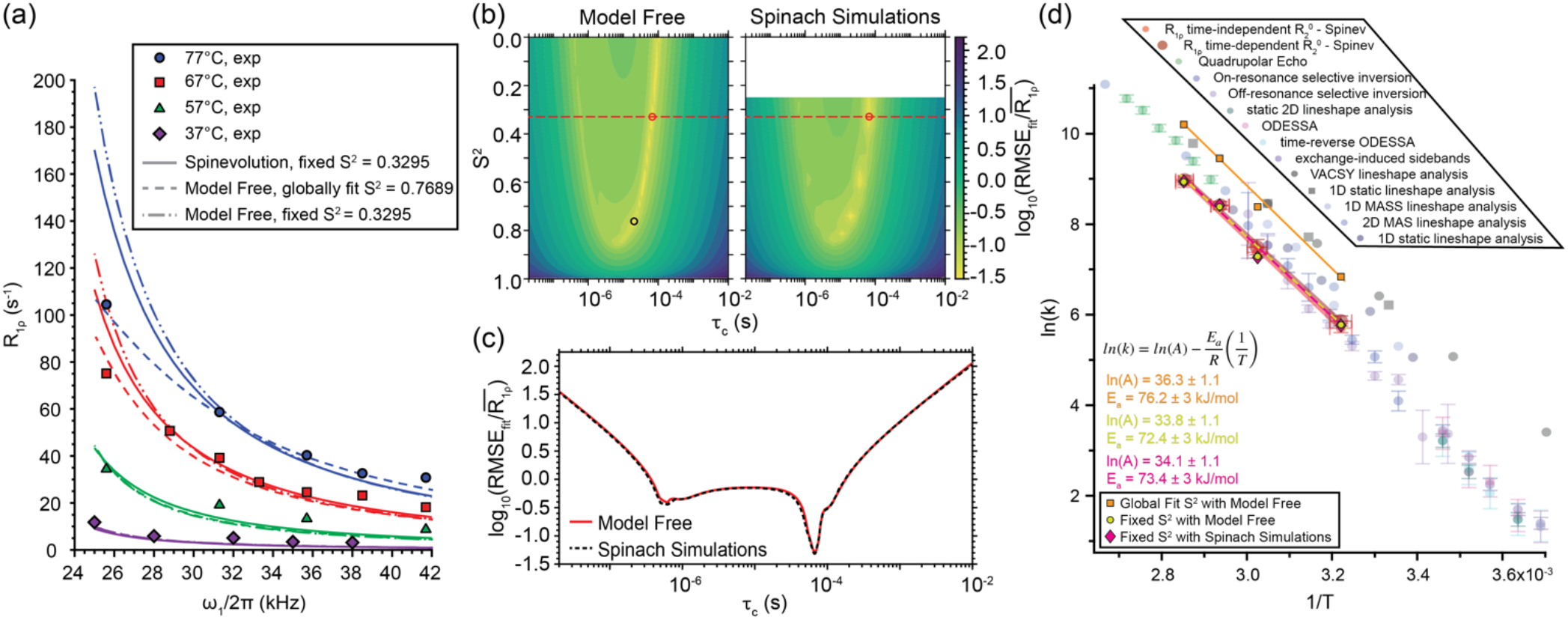
Model free and numerical simulation analysis results for DMS, prior ^42^ and new results with the parameters for the simulations and experiments: ω_oH_/2π = 400 MHz, *v*_r_ = 10 kHz, *v*_1_ = 2.45, 2.75, 3.3, 3.55, 3.8, 4.2 *v*_r_, β = 109° (S^2^ = 0.3295), δ = −36.77 ppm, η = 0 with a temperature independent R_2_^0^ included in the rate constant for each point. **(a)** Experimental (circles), numerical simulation (solid lines), and model free (dashed and dot/dashed lines) relaxation dispersion curves for DMS. **(b)** Contour plots for the normalized RMSE of the 77°C experimental data of the model free and numerical simulations^41^ with the fixed S^2^ best fit (red circle) and global S^2^ best fit (black circle). The residuals for each point were then used to calculate the RMSE_fit_ which was then normalized by the average relaxation rate over the dispersion curve. **(c)** The slice at S^2^ = 0.3295 for the model free and numerical simulations. **(d)** Arrhenius plot for previous studies of DMS by various techniques (Figure 5 from Quinn and McDermott^42^) as well as the three new fits that were evaluated here (orange squares, yellow circles, and pink diamonds) along with their results for ln(A) and E_a_.

In Figure 3(a) experimental relaxation rates as a function of the spin-lock strength ω/2π (aka dispersion curves) are compared to simulations based on different analysis methods used in the previous study and recently. Here, and in general, it may be possible to distinguish broadly between slow, fast, and intermediate exchange.

Proceeding to a quantitative analysis of the data set, it is noteworthy that the basin of acceptable fits shows clear covariance between the timescale and the order parameter. Indeed, one can anticipate that it should be impossible to uniquely determine τ_c_ and S from dispersion curves in either the fast limit or the slow limit kinetic regimes, based on MF expressions. In the fast limit, where k_ex_ >> ω_1_ + ω_r_, we expect a family of equivalently good fits in which S and τ_c_ are covariant, since the dependence of the relaxation rate on τ_c_ and S has the functional form *τ_c_*(1 – *S*)^2^ throughout this kinetic regime (independent of B_0_, ω_1_ or ω_r_), and no manipulation of experimental parameters will allow us to separate or determine unique solutions for S and τ_c_. Similarly, for the slow limit, rates depend on 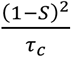 and unique fits are not expected.

Based on the discussion above, one can conclude that prior knowledge of S or the hop angle for the motion is useful determining the timescale in the general case. A global minimum can be identified when the order parameter is fixed to the known value, and is in agreement for the two simulation approaches. For example, for some systems it can be intuited from assumptions about the mechanism of motion in highly symmetric systems. Alternatively, the order parameter can be experimentally determined from measurements of the averaged strength of the interaction that drives relaxation (the CSA or dipolar coupling), and comparison to the static value. For example, measurements of motionally averaged effective dipolar coupling constants have been reported in many cases. ^43 44^ The effect of motion on anisotropic interactions is frequently analyzed with the simplifying assumption of axially symmetric motions, and the effects of motion on the tensor are reported with a single parameter S = 〈*δ*〉/*δ*, where the angle brackets represent the time average. Alternatively, in principle a more nuanced analysis of the effects of motion (order tensor) could be inferred from detailed comparisons of the anisotropy and asymmetry of the tensors at high vs. low temperatures.

DMS is a case in point illustrating the power of restraining S to obtain the timescale of motion. For most of the data the motion is in the slow limit, resulting in a large basin of comparably good fits, whereas assumed prior knowledge of S (or the hop angle and geometry) dramatically narrows the range of compatible timescales (Figure 3(c)). The solid line represents the curves of the best fit solution from numerical simulations as with parameters as provided in Quinn and McDermott,^42^ the dashed line represents the best fit solution of the model free expressions assuming there is one squared global order parameter, S^2^, for the four different temperature points but that this value (S^2^) is not known, and the dot/dashed line is the best fit solution for the points where the S^2^ is fixed at so that S^2^= 0.3295 to correspond with β = 109°. While the dashed line results in a better overall fit for the highest temperature points, it produces an unphysical squared order parameter of 0.7689 which corresponds to an angle difference between the CSA tensors of the two methyl carbons of ~34°. A more detailed investigation of the highest temperature data set (77°C) is shown in Figure 3(b) showing the comparison of the full root mean square error (RMSE) surface of the model free expressions and the numerical simulations. The best fit of the model free expressions to the global model free order parameters is demonstrated with a black circle (global meaning all data sets at all temperatures were fit to the same order parameter). While the model free expressions and numerical simulations produce near identical fitting surfaces with this small CSA system, neither produces a global minimum corresponding to the known order parameter. In other words, if one assumes the hop angle or order parameter to be known (as done previously^42^) the correct rate can be readily identified, but in absence of that knowledge a spurious minimum presents.

The above discussion of the benefit to prior knowledge of the order parameter refers mainly to the fast limit and the slow limit, and not to the intermediate exchange. A unique fit to accurate values for both S and the timescale are potentially possible the intermediate exchange regime. The utility of this fitting procedure to uniquely determine S and τ_c_ is someone hampered by the narrow range of applicable rates in intermediate exchange for which it is successful, making it fortuitous rather than a generally successful approach. Furthermore, peaks in intermediate exchange may be frequently weak, owing in part to the poor transfer efficiency in any pulse sequence that is sensitive to rotating frame relaxation or R2 relaxation. Consequently, the fast limit and slow limit cases discussed above are of keen practical interest.

In many cases the activation energy is of interest and is extracted from rates measured at different temperatures. For example, the energy obtained for DMS from rotating frame relaxation was in excellent agreement with prior reported values (Figure 3(d)). We explored the effect of the covariance of the order parameter and the timescale, on our ability to determine the activation energy. We took the point of view that many motions will have a consistent or similar value for S across a finite temperature range, and that in global fitting S might be constrained to be consistent across temperature. If data were in the fast (or consistently in the slow) limit throughout the temperature range of interest, the effect of constraining S to an erroneous (but consistent) value results in best fit rates that are in error by a consistent multiplicative factor. In this case, when fit to Arrhenius equation, a similar value for Ea will yield. In the example of DMS, activation energies of 73 kJ/mol, 72 kJ/mol, and 76 kJ/mol are obtained. Thus, a relatively consistent activation energy was obtained despite an error in the global order parameter and despite a consistent error in the best fit rates themselves (whereas the entropy of activation, or pre-exponential factor would be expected to be in error).

The reason for a spurious preferred global fit absence of an order parameter restraint may be due to experimental imperfections—that break the degeneracy of the basin of otherwise equivalent fits. For example, artefacts associated with coherent evolution appear in the dispersion curve, and cause elevated relaxation rates in a broad range of applied field strengths near the spinning frequency. These features may result in a better fit with an intermediate exchange small angle motion even if the motion is in the fast or slow regime.

With this in mind, additional simulations illustrate the benefit of using alternative pulse sequences. Simulations for both the traditional spin-lock and RECRR experiments discussed above were tested against model free results. This was accomplished by extracting relaxation rates from a single exponential fit of the numerically simulated decay of magnetization, and then comparing this relaxation rate to the model free expression. Two systems were used, the first was DMS (described above) with five ω_1_ points between 2.45 and 3.8 ω_r_ and the second was an aromatic-like carbon with five ω_1_ points from 0.43 and 2.33 ω_r_ (see Figure 4 for full details). With an assumed spinning frequency of 10 kHz, the methyl group of DMS has an anisotropy smaller than the spinning frequency, resulting in limited coherent evolution of the CSA during the spin-lock. By contrast, for the aromatic carbon there is significant coherent evolution even with the spinning frequency of 16 kHz. Thus, these two cases illustrate two important regimes that may be encountered. Both systems were simulated assuming a range of correlation times (10^-8^ (10 ns) to 10^-3^ s (1 ms)). The simulation assumed hop angles of β = 109° for DMS and 120° for the aromatic carbon. Resulting simulated data were fit both with and without the a priori knowledge of the order parameter. The results of the model free fits to traditional spin-lock experiments for the DMS system are shown in Figure 4(a). The results displayed in figure 4(a) demonstrate that model free expressions can determine the proper correlation time for traditional spin-lock experiments if the order parameter is known a priori and the anisotropy is comparable or smaller than the spinning frequency. Specifically, model free analysis for the methyl group of crystalline DMS yielded rates within 20% of the initial correlation time for most time scales. However, it is noteworthy that the ratio of the fit best τ_c_ to the initial τ_c_ (τ_fit_/τ_in_) deviates significantly from 1 (as does the order parameter) if the order parameter is not assumed to be known, outside of a limited range of τ_c_ (~1 order of magnitude centered around 10^-5^). This success for DMS is principally due to the small anisotropy, namely that it is smaller than the spinning frequency, so that little coherent evolution takes place.

**Figure 4:**
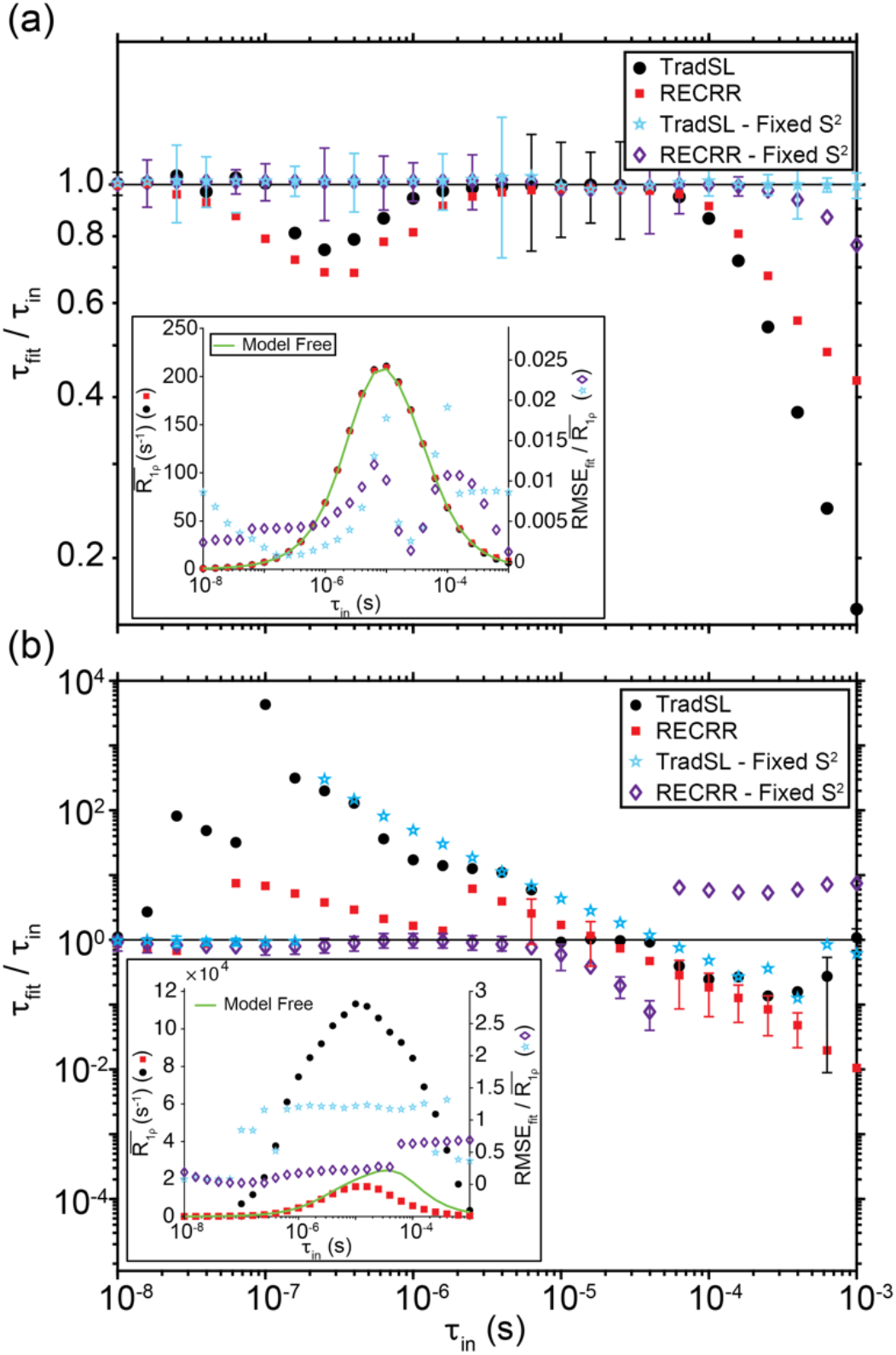
Comparison of the model free analysis best fit correlation time (τ_fit_, y axis value)and the assumed/correct input correlation time of the numerical simulation (τ_in_, x axis value) for DMS **(a)** and an aromatic-like carbon **(b).** Both the traditional spin-lock and RECRR experiments were simulated and the model free fits were performed both with and without the a priori knowledge of S^2^. The inset shows the average relaxation rate (R_1ρ_) for each simulated dispersion curve (averaged over all ω_1_ points on the curve) and from the model free expressions (left axis). The right axis of the inset is the ratio of the best fit RMSE normalized with the average relaxation rate for the simulation. The inset for the aromatic carbon illustrates the better agreement of RECCR with MF expressions. The smallest error for each τ_in_ is shown, errors are considerably larger in some cases. Details of the input system: DMS, ω_oH_/2π = 400 MHz, *v*_r_ = 10 kHz, *v*_1_ = 2.45, 2.75, 3.3, 3.55, 3.8*v*_r_, β = 109° (S^2^ = 0.3295), δ = −36.77 ppm, η = 0 and Aromatic carbon, ω_oH_/2π = 750 MHz, *v*_r_ = 10 kHz, *v*_1_ = 0.43, 0.63, 0.83, 2.16, and 2.33*v*_r_, β = 120° (S^2^ = 0.4395), δ = 175 ppm, η = 0.

Also, it is due to the fact that there is a single dominant relaxation process (no dipolar couplings in this particular system). In this regime, both numerical simulations and model free expressions produce best fits with nearly identical relaxation rate constants, as illustrated in the figure inset. Additionally, for small or moderate anisotropy systems (compared to the spinning frequency), RECRR and the traditional spin-lock result in comparable results.

By contrast, figure 4(b) shows the results of fitting simulated data for an aromatic carbon undergoing ring flip, where an anisotropy that is larger as compared to the spinning frequency is assumed. These results demonstrate that, by contrast to smaller anisotropy systems like the methyl group of DMS (above), the larger anisotropy system produced fits that were frequently several orders of magnitude off from the correct (assumed/simulated) τ_c_ value. Specifically, with the traditional spin-lock, and absent a priori knowledge of the order parameter, the fits gave a unique minimum, but these minima were rarely within 20% of the correct value. For large anisotropy systems (here the aromatic carbon) use of RECRR in place of the traditional spin-lock for the experiments, and a priori knowledge of the order parameter during data analysis, resulted in improved ability to discern the correct timescale with model free analysis. The inset of the figure clearly demonstrates that RECRR outperforms the traditional spin-lock in terms of extending the domain of applicability of the model free formalism for anisotropic systems; the numerical simulations match the model free determined relaxation rate for a range of assumed correlation times, which is particularly evident for fast exchange where τ_c_ < 10^-5^. Overall, even including the improvements from RECRR, the normalized best fit RMSEs are larger than those found for a smaller anisotropy system, and the precision for the derived correlation times correspondingly poorer.

To address the difficulties with model free analysis for anisotropic systems highlighted above, numerical simulations of solid state NMR rotating frame relaxation experiments were also used as a data analysis tool. In this approach, a grid of simulations employing the known spectroscopic parameters, and assuming various motion models is compared with the experimental data. Figure 5 presents RMSE differences between normalized experimental decay constants and simulated decay constants, presented as a heatmap of an unweighted contributions from all points on the dispersion curve. Loci of good agreement between simulation and experiment appear in yellow. An aromatic carbon undergoing two-site flipping is analyzed, with assumed timescales of τ_c_ = 29.5 (a and b) and 790 μs (c). Several conclusions are illustrated by these contour plots. The proper order parameter and timescale corresponds to the global minimum of the numerical simulations for the RECRR experiments (b and c, top panel). This is not the case if model free analysis is used (b and c bottom panel) nor if the traditional spin-lock is used (a). If the order parameter is assumed to be known, the best agreement model free fits nevertheless do not correspond to the correct rate.

**Figure 5:**
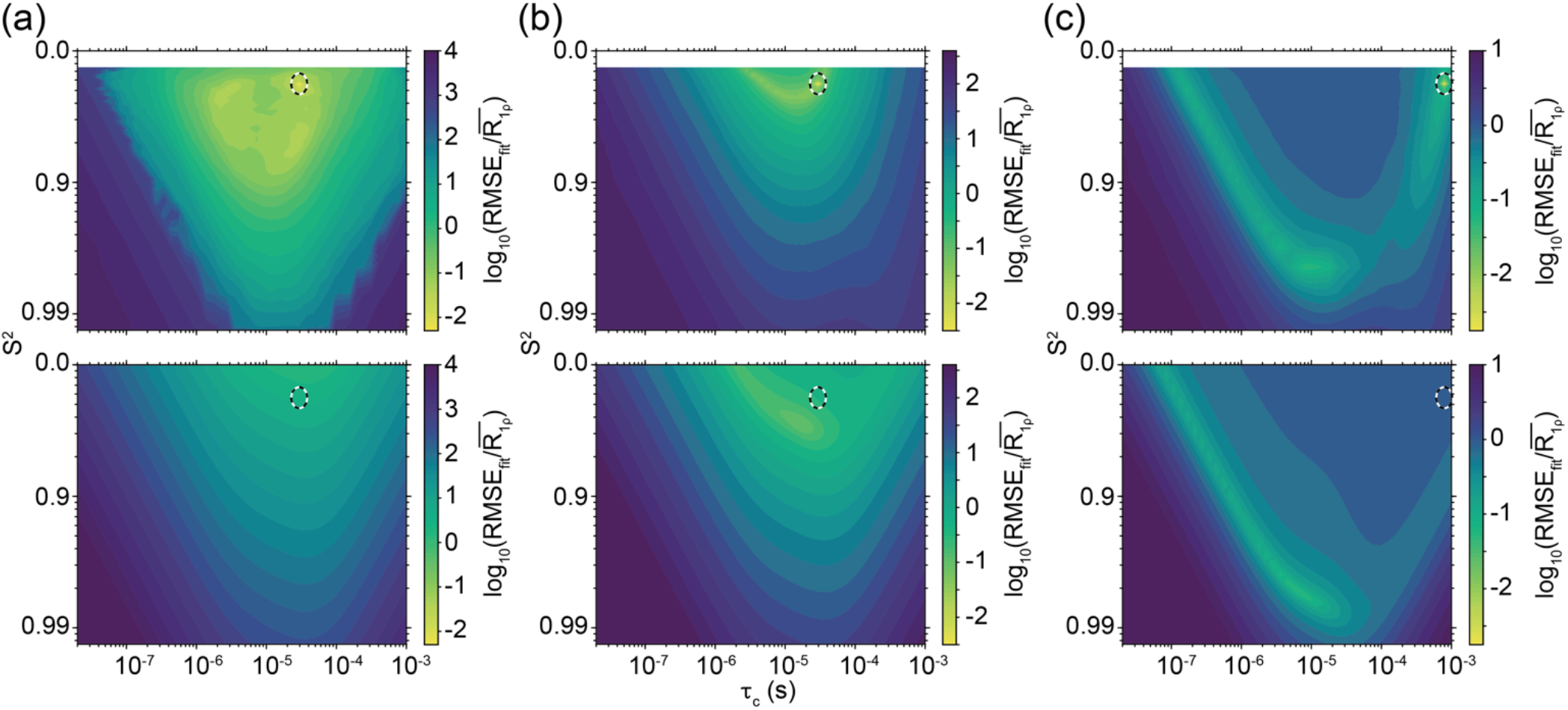
Contour plots of the RMSE comparing a numerically simulated relaxation dispersion curve (CSA mechanism, aromatic carbon), for a traditional Spinlock (a) and RECRR (b,c) experiments. These simulated data were compared to a grid of numerically simulated curves (top) or, separately, to model free expressions (Kurbanov) (bottom). 1000 numerically simulated curves were evaluated with σ = 0.10 Gaussian random noise added to them before the residuals were calculated. The average of the residuals for each point were then used to calculate the RMSE_fit_ which was then normalized by the average relaxation rate over the dispersion curve. The system parameters are: δ_CSA_ = 175 ppm, two-fold hop with β = 120° and τ_c_ = 29.5 (a,b) and 790 (c) μs; dispersion curve points ω_1_ = 0.43, 0.63, 0.83, 2.16, and 2.33 ω_r_ and spinning frequency *v*_r_ = 16 kHz. The black/white dashed oval indicates the correct values used in input parameters for the numerical simulation to create the “test data” relaxation dispersion curve being fit. Note that a more convergent minimum appears with numerical simulations (top) than for model free fitting (bottom). Also significantly narrower basin of acceptable fits are seen for the RECRR experiments (b) vs the TradSL (a). If the RECRR experiment is fit using numerical simulations, the globally best fit returns the correct rate for a broad range of timescales, an aspect under further investigation elsewhere.

Additionally, if the order parameter is assumed to be known, the traditional spin-lock exhibits a broad minimum. These observations require further exploration to see if the RECRR and numerical simulations generally are a reliable approach. Meanwhile, at this stage they provide optimism and motivation for using RECRR or similar experimental strategies and numerical simulations for studies of dynamics using anisotropic systems at moderate spinning frequencies.

Notably, incorrect MF fits for many cases wrongly return high order parameters and intermediate exchange timescales (see figure 5c, bottom for an example). This tendency may result from coherent evolution of the magnetization, which has the effect of broadening the rotary resonance conditions, resulting in dispersion curves that more resemble intermediate exchange timescale model free simulations than fast or slow limit model free simulations. Other effects such as inhomogeneous RF fields may also lead to a similar systematic error.

Additional fit precision can be afforded by simulating the decay curves in the time domain with fitting data to extract a rate constant (Figure 6). The decay curves are not in general strictly exponential, due to coherent evolution and relaxation anisotropic effects, and potentially effects of more complex kinetic models, and therefore avoiding the requirement that the data be fit to an exponential can improve the accuracy and precision of fitting.

**Figure 6:**
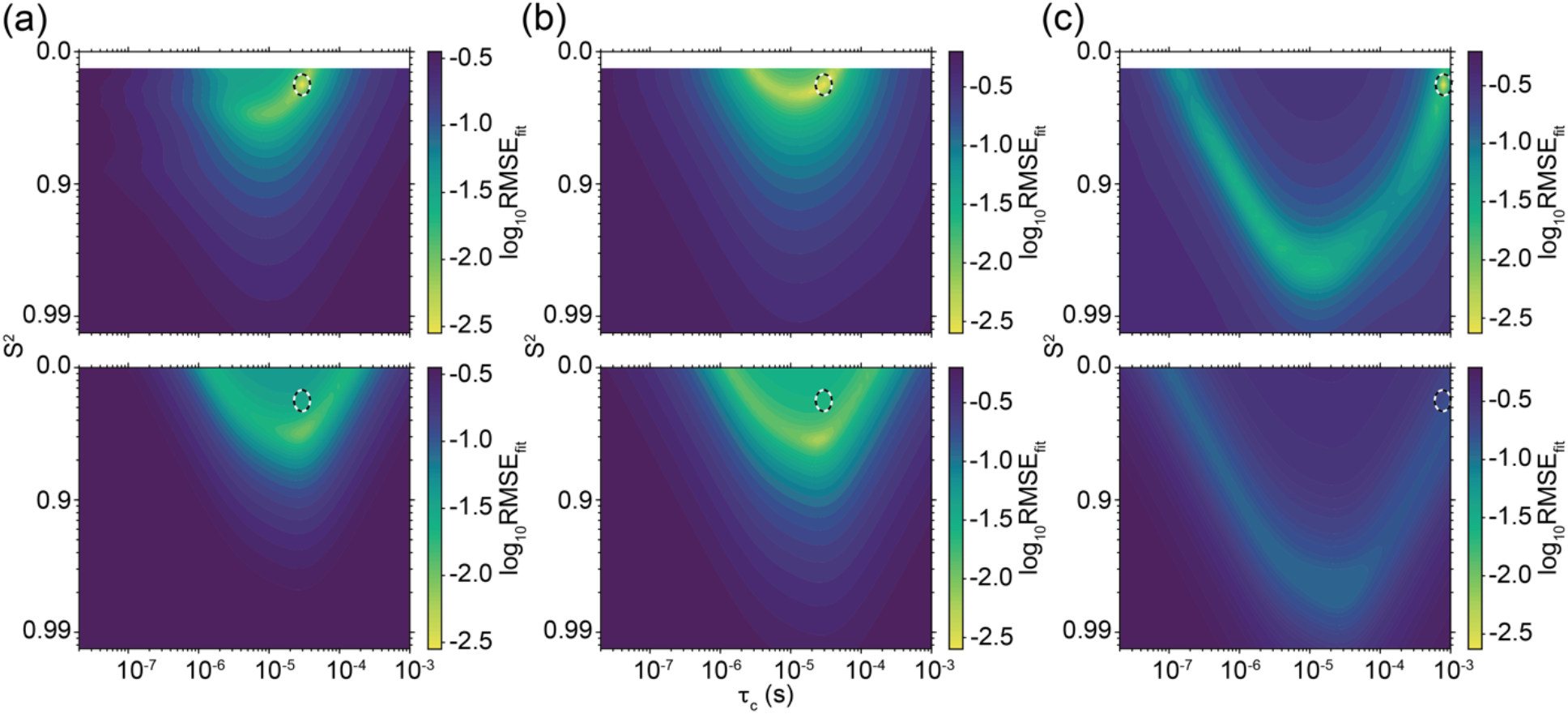
Contour plots of the RMSE comparing the time decays of a numerically simulated relaxation dispersion curve (CSA mechanism, aromatic carbon), for a TradSL (a) and RECRR (b,c) experiments. These simulated data were compared to a grid of decays of numerically simulated curves with 7 points (0, 0.25, 0.75, 1.25, 2.5, 4.375 (4.0, RECRR), 6.25 (6.0, RECRR) ms) (top) or, separately, to decays generated from the single exponential rate of model free expressions (Kurbanov) for the same time points in the numerical simulations (bottom). 1000 numerically simulated curves were evaluated with σ = 0.10 Gaussian random noise added to them before the residuals were calculated. The average of the residuals for each point were then used to calculate the RMSE_fit_. The system parameters are: δ_CSA_ = 175 ppm, two-fold hop with β = 120° and τ_c_ = 29.5 (a,b) and 790 (c) μs; dispersion curve points ω_1_ = 0.43, 0.63, 0.83, 2.16, and 2.33 ω_r_ and spinning frequency *v*_r_ = 16 kHz. The black/white dashed oval indicates the correct values used in input parameters for the numerical simulation to create the “test data” relaxation dispersion curve being fit. Note that a more convergent minimum appears with numerical simulations (top) than for model free fitting (bottom). While, it appears a significantly narrower basin of acceptable fits is present in the TradSL (a) vs the RECRR (b) experiments this is due to the exact correct values of the anisotropic tensor being used in every simulation on the grid allowing the coherent evolution present in the TradSL simulations to provide an additional vector for reducing the RMSE of the fit. If the RECRR experiment is fit using numerical simulations, the globally best fit returns the correct rate for a broad range of timescales, an aspect under further investigation elsewhere.

To summarize, for favorable systems near intermediate exchange (ω_1_ + ω_r_ ~ k_ex_) dispersion data and spinning frequency dependent data can be used to determine both the rate and the hop angle/order parameter. On the other hand, with prior knowledge of the angle or order parameter the time constant can be determined more generally. Lacking prior knowledge of the order parameter, activation energies can nevertheless be accurately obtained.

Faster MAS also presents a strategy for improving fidelity of data, as mentioned above, by removing unwanted coherent evolution. We examined a case of practical interest, the ^13^C carbonyl measured with a MAS frequency of 60 kHz. Analogously to the studies described above, an aspect of interest is whether a dispersion curve (simulated, and therefore with known timescale and angles, and with realistic noise co-added) would have a unique and correct best model free fit. In Figure 7, a carbonyl carbon undergoing two-site flipping is analyzed, with assumed timescale and order parameter of τ_c_ = 90 μs and S^2^ = 0.9918 (6°). This analysis indicates that when evaluating a R_1ρ_ dispersion curve, taken with 5 points below *v*_r_, when utilizing fast spinning on an environment with moderate CSA the TradSL and RECRR experiments are more similar than in the moderate spinning with large CSA case. However, most notably the model free evaluation of the data leads to a strong covariance between the fitted τ_c_ and S^2^ demonstrating a lack of clear minimum if the order parameter is unknown. Additionally, even when the order parameter is known, the minimum fails to fall in the correct position when using the model free expressions. The numerical simulations show a clear minimum at the correct point; however, there are additional minima that appear in covariant line in τ_c_ and S^2^ space (Figure 7, top), but if the order parameter is known prior to fitting the numerical simulation fit yields the expected result.

**Figure 7:**
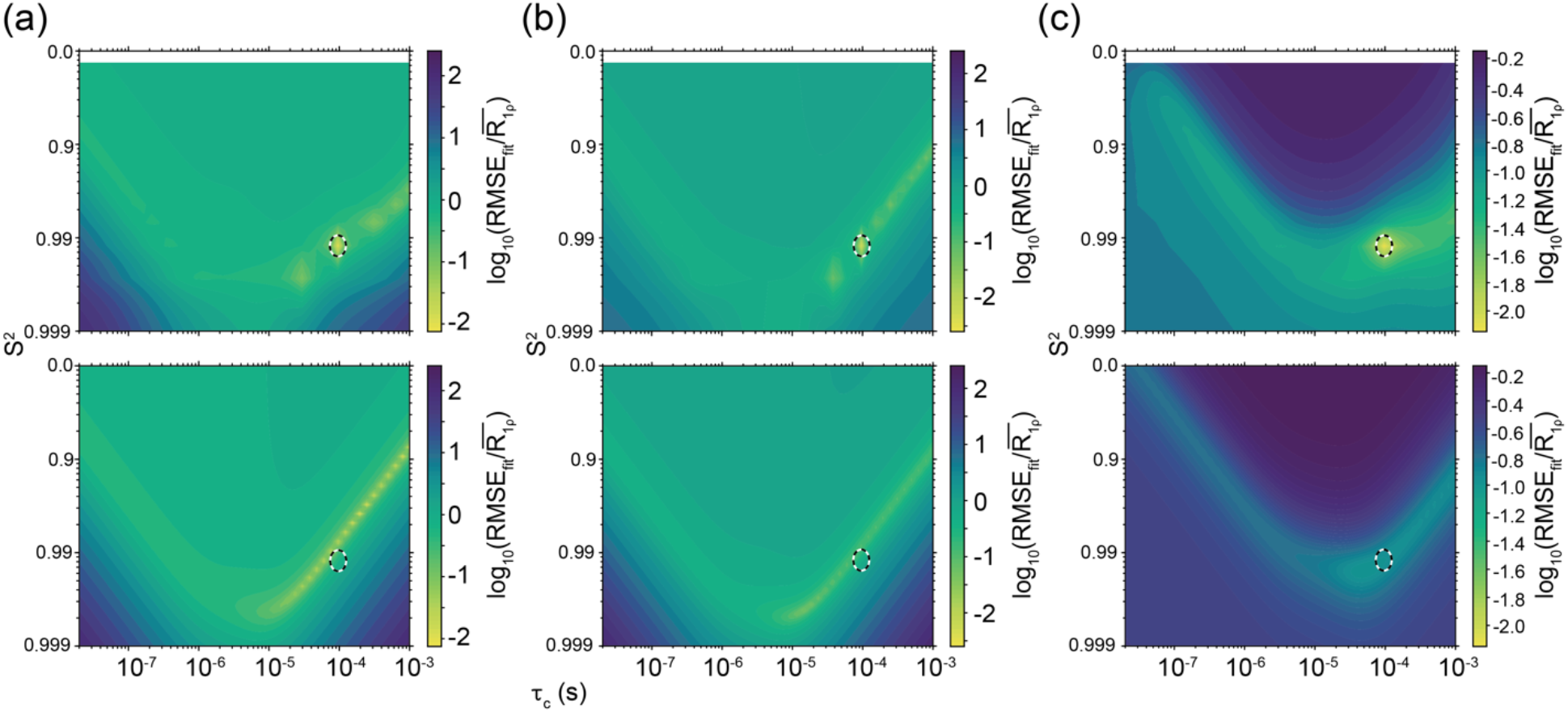
Contour plots of the RMSE comparing a numerically simulated relaxation dispersion curve (CSA mechanism, carbonyl), for a traditional Spinlock (a) and RECRR (b,c) experiments for 900 MHz instrument (^1^H frequency), assuming a magic angle spinning frequency *v*_r_ = 60 kHz, and dispersion curve with *v*_r_ points at 10, 20, 30, 35, and 45 kHz, chosen to keep RF irradiation in realistic ranges and to avoid CSA matching conditions. The assumed values for the correlation time τ_c_ and squared order parameter S^2^ were 90 μs and 0.9918 (6°) and are indicated with a black/white dashed oval. These simulated data were compared to a grid of numerically simulated curves (top) or, separately, to model free expressions (Kurbanov) (bottom). The data in (c) was compared to the time decays instead of the fitted relaxation rates. 500 numerically simulated curves were evaluated with σ = 0.10 Gaussian random noise coadded before calculating residuals. A unique and correct fit can be obtained using numerical simulations, without prior knowledge of the angle or order parameter whereas the fits via model free are incorrect and not unique due to the covariance of the fitting parameters. The minimum is particularly robust when using the time domain decay points in the place of the extracted relaxation rates. For the numerical simulation the CSA tensor is defined as: δ = −90 ppm and η = 0.6 and for the model free the CSA tensor is defined as: δ = −90 ppm and η = 0.

Overall, the discussion above indicates some hope for extracting quantitative descriptions of motion from SSNMR rotating frame relaxation studies, with a number of important caveats highlighted, including suppression of coherent evolution, the use of appropriate simulation strategies and in many cases prior estimates of the angle or order parameter. In this discussion, numerous simplifications were also imposed. Our analyses were restricted to an assumed single motion or correlation time. In many systems, more complex motions might be encountered and fitting these motions will presumably require larger sets of data. Given the complexity of biopolymer motions, identifying strategies for correct and accurate decomposition of multiple motions represents an important challenge for the future. Looking to the future, we face the exciting challenge of harnessing the many new technical developments in solid state NMR to enables study complex dynamics.

## Concluding Remarks

In light of the burgeoning number of applications of rotating frame relaxation in magic angle spinning solid state NMR experiments, we expect that there is a bright future for this area of spectroscopy. We explored approaches to analysis, focusing on systems with simple kinetics of exchange (two-site hop) where the chemical shift anisotropy provides the dominant mechanism. We conclude that a correct and precise timescale can be obtained if a number of criteria are met. Beyond the need for high signal-to-noise decay profiles for resolved and assigned peaks, we emphasized the need for suppression of coherent evolution. For this reason, for systems with large CSA, pulse sequences that suppress coherent evolution or ultrafast spinning is needed. If coherent evolution is not fully suppressed, numerical simulations may be required to obtain confident and correct information about the motions. When all of these criteria are met, for favorable systems near intermediate exchange (w_1_ + w_r_ ~ k_ex_) dispersion (w_1_ dependent) data can be used to determine the timescale of motion. More generally, considering fast and slow limit cases, prior knowledge of the angle or order parameter is needed to determine the time constant with confidence. This is illustrated with a case with relatively narrow shielding anisotropy compared to the spinning frequency, (a methyl group), where model free based fitting protocols identified correct rate constants if S is constrained by knowledge of the molecular geometry (angles) of the motion. In absence of prior knowledge covariance of τ_c_ and S leads to numerous minima in data fitting. Nevertheless, despite the fact that the unique and correct fit cannot be obtained without prior knowledge of the order parameter, the activation energy can be obtained without any prior knowledge in many cases. Going forward, given the powerful developments in faster MAS spinning frequencies and other new technologies, there are many opportunities for this area of science.

## Notes

### Competing Interest Statement

The authors have declared no competing interest.

## References

1. Schanda, P. & Ernst, M. Studying dynamics by magic-angle spinning solid-state NMR spectroscopy: Principles and applications to biomolecules. Prog. Nucl. Magn. Reson. Spectrosc. 96, 1–46 (2016).

2. Rovo, P. & Linser, R. Microsecond Timescale Protein Dynamics: a Combined Solid-State NMR Approach. Chemphyschem 19, 34–39 (2018).

3. Palmer, A. G., Williams, J. & McDermott, A. Nuclear magnetic resonance studies of biopolymer dynamics. J. Phys. Chem. 100, 13293–13310 (1996).

4. Palmer, A. G. & Massi, F. Characterization of the dynamics of biomacromolecules using rotating-frame spin relaxation NMR spectroscopy. Chem. Rev. 106, 1700–1719 (2006).

5. Rovó, P. & Linser, R. Microsecond Time Scale Proton Rotating-Frame Relaxation under Magic Angle Spinning. J. Phys. Chem. B 121, 6117–6130 (2017).

6. Smith, A. A., Testori, E., Cadalbert, R., Meier, B. H. & Ernst, M. Characterization of fibril dynamics on three timescales by solid-state NMR. J. Biomol. NMR 65, 171–191 (2016).

7. Helmus, J. J., Surewicz, K., Surewicz, W. K. & Jaroniec, C. P. Conformational flexibility of Y145stop human prion protein amyloid fibrils probed by solid-state nuclear magnetic resonance spectroscopy. J. Am. Chem. Soc. 132, 2393–2403 (2010).

8. Gauto, D. F. et al. Aromatic ring dynamics, thermal activation and transient conformations of a 468 kDa enzyme by specific labeling and fast-MAS NMR. J. Am. Chem. Soc. jacs.9b04219 (2019). doi:10.1021/jacs.9b04219

9. Öster, C., Kosol, S. & Lewandowski, J. R. Quantifying Microsecond Exchange in Large Protein Complexes with Accelerated Relaxation Dispersion Experiments in the Solid State. Sci. Rep. 9, (2019).

10. Vasa, S. K., Singh, H., Rovó, P. & Linser, R. Dynamics and Interactions of a 29 kDa Human Enzyme Studied by Solid-State NMR. J. Phys. Chem. Lett. 9, 1307–1311 (2018).

11. Lakomek, N.-A. et al. Microsecond Dynamics in Ubiquitin Probed by Solid-State 15 N NMR Spectroscopy R 1ρ Relaxation Experiments under Fast MAS (60-110 kHz). Chem. - A Eur. J. 23, 9425–9433 (2017).

12. Good, D., Pham, C., Jagas, J., Lewandowski, J. R. & Ladizhansky, V. Solid-state NMR provides evidence for small-amplitude slow domain motions in a multi-spanning transmembrane α-helical protein. J. Am. Chem. Soc. jacs.7b03974 (2017). doi:10.1021/jacs.7b03974

13. Jekhmane, S. et al. Shifts in the selectivity filter dynamics cause modal gating in K + channels. Nat. Commun. 10, 1–12 (2019).

14. Krushelnitsky, A., Zinkevich, T., Reif, B. & Saalwächter, K. Slow motions in microcrystalline proteins as observed by MAS-dependent 15N rotating-frame NMR relaxation. J. Magn. Reson. 248, 8–12 (2014).

15. Ma, P. et al. Probing transient conformational states of proteins by solid-state R1ρ relaxation-dispersion NMR spectroscopy. Angew. Chemie - Int. Ed. 53, 4312–4317 (2014).

16. Good, D. B. et al. Conformational dynamics of a seven transmembrane helical protein Anabaena Sensory Rhodopsin probed by solid-state NMR. J. Am. Chem. Soc. 136, 2833–2842 (2014).

17. Lamley, J. M., Öster, C., Stevens, R. A. R. A. & Lewandowski, J. R. Intermolecular Interactions and Protein Dynamics by Solid-State NMR Spectroscopy. Angew. Chemie Int. Ed. 54, 15374–15378 (2015).

18. Lamley, J. M. et al. Unraveling the complexity of protein backbone dynamics with combined 13C and 15N solid-state NMR relaxation measurements. Phys. Chem. Chem. Phys. (2015). doi:10.1039/c5cp03484a

19. Medeiros-Silva, J. et al. 1H-Detected Solid-State NMR Studies of Water-Inaccessible Proteins In Vitro and In Situ. Angew. Chemie - Int. Ed. 55, 13606–13610 (2016).

20. Saurel, O. et al. Local and global dynamics in Klebsiella pneumoniae outer membrane protein A in lipid bilayers probed at atomic resolution. J. Am. Chem. Soc. (2017). doi:10.1021/jacs.6b11565

21. Kurauskas, V. et al. Slow conformational exchange and overall rocking motion in ubiquitin protein crystals. Nat. Commun. (2017). doi:10.1038/s41467-017-00165-8

22. Gauto, D. F. et al. Protein conformational dynamics studied by15N and 1H R1ρrelaxation dispersion: Application to wild-type and G53A ubiquitin crystals. Solid State Nucl. Magn. Reson. (2017). doi:10.1016/j.ssnmr.2017.04.002

23. Shannon, M. et al. Conformational Dynamics in the Core of Human Y145Stop Prion Protein Amyloid Probed by Relaxation Dispersion NMR. ChemPhysChem (2018). doi:10.1002/cphc.201800779

24. Lewandowski, J. R., Sass, H. J., Grzesiek, S., Blackledge, M. & Emsley, L. Site-specific measurement of slow motions in proteins. J. Am. Chem. Soc. 133, 16762–16765 (2011).

25. Kurauskas, V. et al. Cross-Correlated Relaxation of Dipolar Coupling and Chemical-Shift Anisotropy in Magic-Angle Spinning R1 NMR Measurements: Application to Protein Backbone Dynamics Measurements. J. Phys. Chem. B 120, 8905–8913 (2016).

26. Singh, H. et al. Fast Microsecond Dynamics of the Protein-Water Network in the Active Site of Human Carbonic Anhydrase II Studied by Solid-State NMR Spectroscopy. J. Am. Chem. Soc. 141, 19276–19288 (2019).

27. Rovó, P. et al. Mechanistic Insights into Microsecond Time-Scale Motion of Solid Proteins Using Complementary 15 N and 1 H Relaxation Dispersion Techniques. J. Am. Chem. Soc. 141, 858–869 (2019).

28. Jirasko, V. et al. Proton-Detected Solid-State NMR of the Cell-Free Synthesized α-Helical Transmembrane Protein NS4B from Hepatitis C Virus. ChemBioChem 21, 1453–1460 (2020).

29. Tognetti, J., Trent Franks, W., Gallo, A. & Lewandowski, J. R. Accelerating 15N and 13C R1 and R1ρ relaxation measurements by multiple pathway solid-state NMR experiments. J. Magn. Reson. 107049 (2021). doi:10.1016/j.jmr.2021.107049

30. Krushelnitsky, A., Reichert, D. & Saalwächter, K. Solid-state NMR approaches to internal dynamics of proteins: From picoseconds to microseconds and seconds. Acc. Chem. Res. 46, 2028–2036 (2013).

31. Zhehong, G., Grant, D. M. & Ernst, R. R. NMR chemical shift anisotropy measurements by RF driven rotary resonance. Chem. Phys. Lett. 254, 349–357 (1996).

32. Rovó, P. Recent advances in solid-state relaxation dispersion techniques. Solid State Nucl. Magn. Reson. 108, (2020).

33. Keeler, E. G., Fritzsching, K. J. & McDermott, A. E. Refocusing CSA during magic angle spinning rotating-frame relaxation experiments. J. Magn. Reson. 296, 130–137 (2018).

34. Long, J. R., Sun, B. Q., Bowen, A. & Griffin, R. G. Molecular Dynamics and Solid State NMR. J. Am. Chem. Soc. 116, 11950–11956 (1994).

35. Lipari, G. & Szabo, A. MODEL-FREE APPROACH TO THE INTERPRETATION OF NUCLEAR MAGNETIC-RESONANCE RELAXATION IN MACROMOLECULES. 1. THEORY AND RANGE OF VALIDITY. J. Am. Chem. Soc. 104, 4546–4559 (1982).

36. Lipari, G. & Szabo, A. A Model-free approach to the interpretation of nuclear magnetic resonance relaxation in macromolecules. 1. Therory and range of validity. J Am Chem Soc 104, 4546–4559 (1982).

37. Beckwith, M. A., Erazo-Colon, T. & Johnson, B. A. RING NMR dynamics: software for analysis of multiple NMR relaxation experiments. J. Biomol. NMR 75, 9–23 (2021).

38. Kurbanov, R., Zinkevich, T. & Krushelnitsky, A. The nuclear magnetic resonance relaxation data analysis in solids: General R1R1 equations and the model-free approach. J. Chem. Phys. 135, 184104 (2011).

39. Schanda, P. Solid-state NMR studies of protein dynamics: New approaches and applications to crystalline proteins and large molecular assemblies. (2015).

40. Veshtort, M. & Griffin, R. SPINEVOLUTION: A powerful tool for the simulation of solid and liquid state NMR experiments. J. Magn. Reson. 178, 248–282 (2006).

41. Hogben, H. J., Krzystyniak, M., Charnock, G. T. P., Hore, P. J. & Kuprov, I. Spinach - A software library for simulation of spin dynamics in large spin systems. J. Magn. Reson. 208, 179–194 (2011).

42. Quinn, C. M. & McDermott, A. E. Quantifying conformational dynamics using solidstate R1ρ experiments. J. Magn. Reson. 222, 1–7 (2012).

43. Hong, M., Yao, X. L., Jakes, K. & Huster, D. Investigation of molecular motions by Lee-Goldburg cross-polarization NMR Spectroscopy. J. Phys. Chem. B 106, 7355–7364 (2002).

44. Chevelkov, V., Fink, U. & Reif, B. Accurate determination of order parameters from1H,15N dipolar couplings in MAS solid-state NMR experiments. J. Am. Chem. Soc. 131, 14018–14022 (2009).

